# Quantitating denaturation by formic acid: Imperfect repeats are essential to the stability of the functional amyloid protein FapC

**DOI:** 10.1101/2020.03.09.983882

**Authors:** Line Friis Bakmann Christensen, Jan Stanislaw Nowak, Thorbjørn Vincent Sønderby, Signe Andrea Frank, Daniel Erik Otzen

## Abstract

Bacterial functional amyloids are evolutionarily optimized to aggregate to help them fulfil their biological functions, *e.g.* to provide mechanical stability to biofilm. Amyloid is formed in *Pseudomonas* sp. by the protein FapC which contains 3 imperfect repeats connected by long linkers. Stepwise removal of these repeats slows down aggregation and increases the propensity of amyloids to fragment during the fibrillation process, but how these mechanistic properties link to fibril stability is unclear. Here we address this question. The extreme robustness of functional amyloid makes them resistant to conventional chemical denaturants, but they dissolve in formic acid (FA) at high concentrations. To quantify this, we first measured the denaturing potency of FA using 3 small acid-resistant proteins (S6, lysozyme and ubiquitin). This revealed a linear relationship between [FA] and the free energy of unfolding with a slope of *m*_FA_, as well as a robust correlation between protein residue size and *m*_FA_. We then measured the solubilisation of fibrils formed from different FapC variants (with varying number of repeats) as a function of [FA]. The resulting *m*_FA_ values revealed a decline in the number of residues driving amyloid formation when at least 2 repeats were deleted. The midpoint of denaturation declined monotonically with progressive removal of repeats and correlated with solubility in SDS. Complete removal of all repeats led to fibrils which were solubilized at FA concentrations 2-3 orders of magnitude lower than the repeat-containing variants, showing that at least one imperfect repeat is required for the stability of functional amyloid.

## INTRODUCTION

The term ‘amyloid’ is normally associated with misfolding of different proteins, resulting in neurodegenerative diseases like Alzheimer’s and Parkinson’s disease, but the number of cases where the amyloid structure is used for functional purposes is steadily increasing^*1, 2*^. The first functional amyloid to be identified and purified were the curli fibrils expressed by *E. coli* and *Salmonella enteritidis*^*3, 4*^. Since their discovery, curli has been shown to be widely expressed among different bacteria, spanning at least four different phyla^*5*^. Curli fibrils serve an architectural role in bacterial biofilm but are also involved in cell attachment and invasion^*6-8*^. Curli are composed primarily of the protein CsgA which, for bacterial strains within the Enterobacteriales and the Vibrionales, is expressed from the *csgBAC* operon together with the nucleator protein CsgB and the chaperone CsgC^*5, 9, 10*^. Furthermore, four additional Csg proteins are expressed from the *csgDEFG* operon and they act as a transcription regulator of the *csgBAC* operon (CsgD), chaperones (CsgE/CsgF) and as an outer membrane pore protein (CsgG)^*9-11*^. All Csg proteins, except CsgD, are targeted for Sec-dependent secretion across the inner bacterial membrane to the periplasm^*12*^.

A decade ago, we identified a similar amyloid system in *Pseudomonas* termed *f*unctional *a*myloid in *Pseudomonas* (fap)^*13*^. The fap system encodes proteins FapA-F expressed from a single *fapA-F* operon. Like the curli system, it encodes an outer membrane pore protein (FapF) and possible chaperones (FapA and FapD) besides the primary amyloid-forming protein FapC and a potential nucleator FapB^*14*^ (Fig. 1). Similar to the curli system, fap fibrils contribute to biofilm formation^*15, 16*^ and play roles in virulence^*17*^ and the individual Fap proteins are also secreted across the inner membrane via Sec^*15, 18, 19*^. However, the fap system is evolutionarily younger than the curli system and only exists within a single phylum, the Proteobacteria^*20*^.

**Figure 1.**
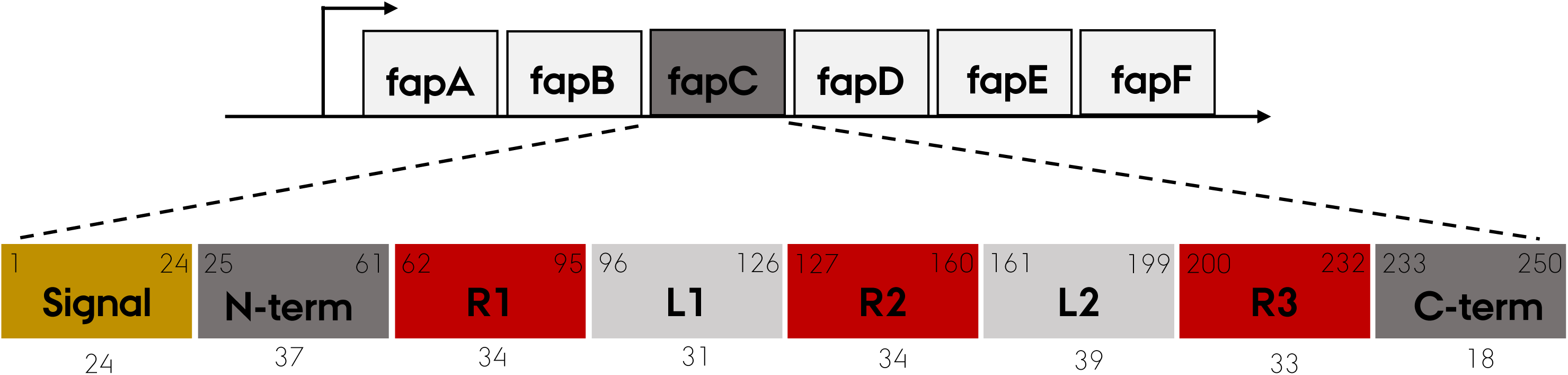
The *fapA-F* operon and the FapC protein. The *fapA-F* operon encodes the six Fap proteins, FapA-F. The *Pseudomonas* sp. UK4 FapC protein consists of a 24 aa N-terminal signal peptide (yellow) and three imperfect repeats (R1—R3, red) separated by two linker regions (L1-L2, light grey). Start and end amino acid positions for the different regions are shown in the upper corners and the length is shown below.

Both amyloid proteins CsgA and FapC consist of multiple imperfect repeats. CsgA has five, each ca. 20 residues in length and folded as individual β-hairpins separated by a tight turn (4-5 residues) according to a computationally predicted structure^*21*^. In support of this, peptides corresponding to three of the individual repeats (repeat 1, 3 and 5) readily form amyloid on their own^*22*^. FapC has 3 longer repeats (ca. 35 residues long) which are also predicted to form β-hairpins^*23*^. However, in FapC the repeats are separated by linker regions of variable length; especially the second linker shows large variations in size with lengths ranging from 39 (in the *Pseudomonas sp.* UK4 strain used in this study) to more than 250 residues for *Pseudomonas putida* F1^*13*^. For FapC, stepwise removal of these repeats has only relatively little effect on the rapidity of fibrillation but increases the ensuing fibrils’ tendency to fragment during shaking, thus forming new growing fibrils^*24*^. While that study focused on the *mechanistic* roles of the different repeats, we here address the question of how removal of these repeats affects the *stability* of the formed fibrils. First of all, this requires a reliable assay to determine fibril stability.

For globular monomeric proteins, stability is defined by the distribution between native and denatured states. Analogously, there is an equilibrium between the fibrillated and monomeric state; the higher the concentration of the monomeric species (corresponding to a *c*ritical *a*ggregation *c*oncentration, *cac*), the less stable the fibril. Consequently, the free energy of fibrillation can be expressed as *–RT* ln *cac*^*25*^, and can be further analysed by displacing the equilibrium towards the monomeric state by *e.g.* addition of denaturants^*26*^. This formalism has *i.a.* been used to evaluate the stability of the Aβ peptide, aided by the relatively high Aβ *cac*^*25*^. However, fibrillation of functional amyloids such as FapC is very efficient and typically leads to impracticably low levels of monomer^*13*^, even in the presence of high concentrations of denaturant (data not shown) or boiling sodium dodecyl sulfate (SDS)^*13*^. Conveniently, high concentrations (typically > 80%) of formic acid (FA) have been shown to solubilize functional amyloids. This phenomenon even serves as an operational basis to identify functional amyloid in complex mixtures^*27*^. The extent to which a protein fibril is dissolved by a progressive increase in FA may therefore provide a measure of the fibril’s stability. To investigate this systematically, however, we need a more general analysis of the denaturation potency of FA. To the best of our knowledge, such an analysis has not been performed yet and we here provide it as part of our analysis of the stability of functional amyloid.

Formic acid – HC(O)OH – is the simplest carboxylic acid. As a protein solvent, FA is superior to most common organic solvents (*e.g.* glycerol, DMSO or trifluoroacetic acid), solubilizing the protein polypeptide chain through protonation, destabilization of hydrogen bonds, and hydrophobic residue interactions^*28, 29*^. High concentrations (> 70% v/v) of FA solubilize large, fibrous protein complexes such as collagen, wool keratin, and silk fibroin^*30-32*^; the FA-solubilized form of fibroin seem to be more compact than when solubilized in water, although fibroin is in random coil conformation in both solutions^*30*^. Solubilization does not involve chemical modification^*30*^; FA can formylate Thr and Ser (*O*-formylation) as well as Lys (*N*-formylation), but this requires high concentrations (> 80% v/v) and extended time intervals (many hours)^*28*^. The amide derivative of FA, formamide (FM), is also able to solubilize and unfold globular proteins such as myoglobin, cytochrome c and insulin^*33, 34*^. Though not an acid, it still interacts with proteins through hydrogen bond disruption and solubilization of hydrophobic surfaces^*35*^. Thus, FM constitutes a protonation-free comparison to FA.

Our objectives for this study are two-fold: (1) to establish a quantitative basis for the use of FA to determine protein stability and (2) to employ this approach to determine the contribution of the individual repeats in FapC to fibril stability. To achieve goal 1, we investigate the impact of FA on the stability of proteins which unfold according to a simple two-state system, making it straightforward to assess the impact of FA on their stability. However, given FA’s properties as acid, the simple lowering of pH will in itself make a major contribution to FA’s destabilizing properties. To extract the non-protonation-related denaturing properties of FA we need to be able to subtract the contributions that a simple drop in pH would make to destabilization. This restricts our investigations to proteins which do not unfold at low pH unless other denaturing measures are involved, such as heat. Therefore we have selected three acid-resistant proteins (hen egg white lysozyme^*36-39*^, S6^*40-43*^ from *Thermus thermophilus* and bovine ubiquitin^*44-50*^) whose stabilities have all been extensively analyzed under a range of conditions. For all three proteins, we determine their stability in FA based on near-UV circular dichroism thermal scans over a range of FA concentrations. For each FA concentration, we carry out a thermal scan at the corresponding pH adjusted with HCl and subtract this effect from that of FA to quantitate the intrinsic contribution of FA to protein stability. Furthermore, we carry out these experiments using < 30% v/v FA to minimize formylation^*28*^. To achieve goal 2, we use the obtained relationship between FA concentration and its denaturation potency (*m*-values) to evaluate the FA depolymerization assays of eight different FapC constructs and thereby quantify the aggregative effect of individual repeats of FapC.

## RESULTS AND DISCUSSION

### Near-UV CD reveals reversible and cooperative thermal unfolding of acid-stable proteins in formic acid despite local relaxation of the native structure of S6 and Ubi

To quantitate the denaturing potency of formic acid (FA), we used near-UV circular dichroism (CD) to carry out thermal scans of three model proteins in different concentrations of FA. The proteins are the 129-residue hen egg white lysozyme (HEWL), the 101-residue ribosomal protein S6 from *Thermus thermophiles* and the 76-residue bovine ubiquitin (Ubi). These proteins were selected for a number of reasons. Firstly, they were well-characterized small proteins in a range of sizes (going from 8.6 kDa through 12 kDa to 16.2 kDa) which have previously been used as model systems for fundamental studies in protein stability and folding^*36-50*^. Secondly, they unfold in a single step without equilibrium intermediates. Thirdly, they remain folded in acidic conditions, allowing us to carry out thermal scans at low pH to determine melting temperatures in the presence of FA. The protein solutions were not buffered beforehand, so addition of even small amounts of FA led to a large drop in pH. Due to the high absorption of FA in the far-UV range, we monitored protein unfolding using near-UV CD. Near-UV CD is highly sensitive to correct folding of the protein; indeed, it can be just as sensitive to loss of native structure as enzyme activity^*51*^. Before we proceed further, we will therefore describe the impact of FA on the structures of our 3 model proteins as indicated by near-UV.

The near-UV signal of lysozyme in FA at room temperature (RT) is essentially identical to that in pure water (Fig. S2AB), showing that the protein remains natively folded in FA. In the case of S6, there is a 4-5 fold reduction in signal intensity in FA at RT compared to that in water, but there is still a distinct spectroscopic signature centered around 280 nm which disappears at 90°C and is largely regained upon cooling down (Fig. S2CD). We have previously observed that S6 adopts a more flexible or “quasi-native” (but not molten globule-like)^*52*^ structure at low pH. The single Trp residue in S6 (position 62) hydrogen bonds to Glu5 and Glu41 in the crystal structure of S6 (PDB: 1RIS^*53*^), and these residues are likely to be protonated at low pH. This will most likely disrupt hydrogen bonding with Trp and increase its mobility, leading to a reduction in near-UV CD. In 10 mM HCl (pH 2) S6 still folds and unfolds according to a two-state transition with well-defined kinetics^*52*^, but the change in compactness upon unfolding (as determined by the so-called *m*-value, which describes how the free energy of unfolding changes with denaturant concentration) is reduced by ca. 40%. This is most likely due to an expansion of the native state, as indicated by the loss in near-UV signal combined with an increased affinity for the fluorescent probe ANS, which preferably binds to exposed hydrophobic patches, as well as decreased sensitivity of this “low-pH” state to mutagenesis^*52*^. As regards Ubi, there is a shift in the relative intensities of the 3 peaks seen in the near-UV CD signal at 265, 275 and 281 nm (Fig. S2EF). In water, these peaks all rise above the downward turn seen from ca. 295 nm down to 287 nm to reach positive ellipticity values. In FA, they are still visible but do not rise so much, suggesting a less asymmetric (rigid) environment for the aromatic groups at low pH. The spectrum is similar to the structure of the V62A mutant of ubiquitin assumed at pH 2-4, which according to NMR retains native secondary and tertiary structure, except for the unfolding of a long loop between two β-strands^*46*^. Thus, both S6 and Ubi retain large parts of their original structure and overall compactness in FA but undergo some loosening of the tertiary fold.

Despite S6’ and Ubi’s spectroscopic changes, all three proteins unfold reversibly and cooperatively in thermal scans, leading to a sigmoidal denaturation curve which can be fitted to a two-state unfolding transition from the (quasi-)native state N to the denatured state D (Fig. 2). Furthermore, the refolded protein curves retain most of the signal after cooling the protein back to RT (Fig. S2A,C,E). This justifies the use of a reversible unfolding model to fit the data. The fits provide both the temperature midpoint of denaturation, *T*_m_, and the enthalpy of denaturation at *T*_m_, ΔH_*T*m_.

**Figure 2.**
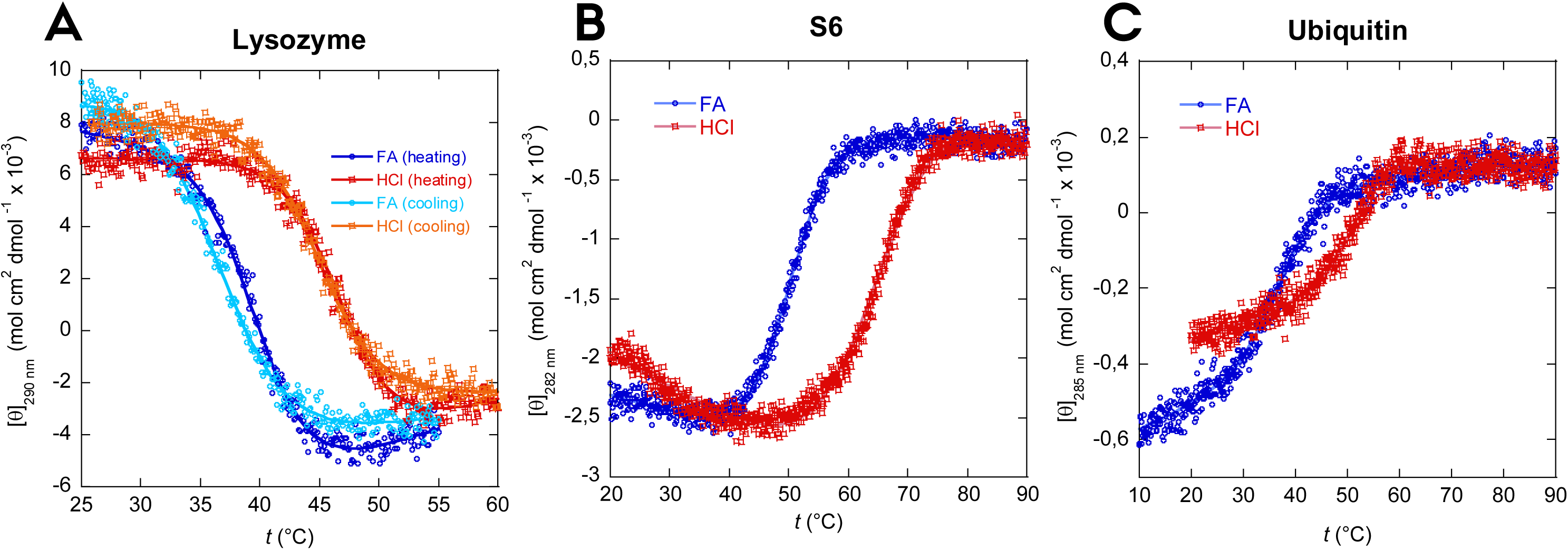
Thermal unfolding curves of (A) lysozyme, (B) S6 and (C) ubiquitin in 10% v/v FA and in HCl solutions with the corresponding pH. In 10% FA all three proteins show significantly reduced thermal stability while maintaining a reversible two-state unfolding (Figure S2). The sigmoidal curves were fitted using eq. (1).

### Formic acid reduces T_m_ more than HCl at similar pH values

Thermal scans were performed on protein solutions containing 0.1% - 25% v/v FA (equal to 0.27-6.63 M given FA’s density of 1.22 g/mL and molecular weight of 46 Da in the protonated form). All three proteins show a linear decrease in *T*_m_ (and thus *t*_m_) with FA concentration (Fig. 3). To separate effects of low pH from those of the FA molecule, we carried out thermal scans in HCl at the same pH values as those obtained with FA. At low FA concentrations, *t*_m_ values of unfolding in FA follow those in HCl (Fig. 4) indicating that denaturation at those concentrations is a simple pH effect. However, as FA rises above 0.5 M, the pH effect caused by HCl starts to level out (*t*_m_ remains essentially constant for S6 and Ubi at all pH values and levels out for HEWL around pH 1.4 which corresponds to a FA concentration of 3 M), while the denaturation effect of FA continues unabated.

**Figure 3.**
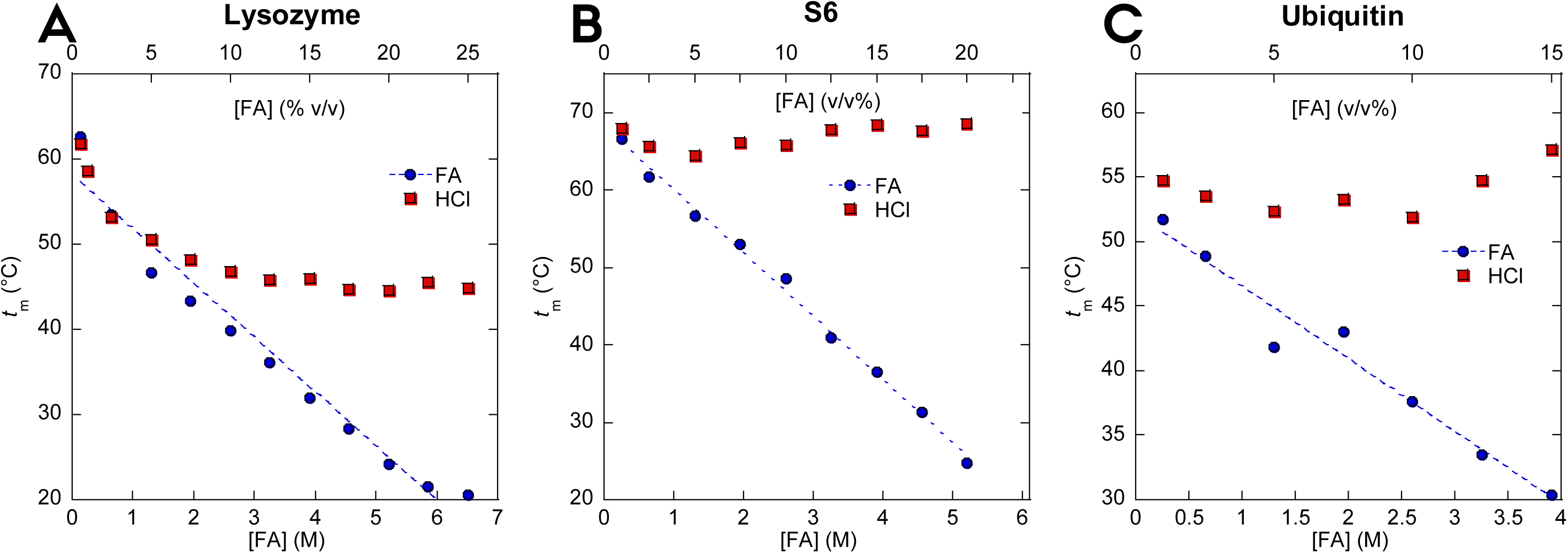
Melting temperature of (A) lysozyme, (B) S6 and (C) ubiquitin. The FA concentration is represented both as molar concentration (M) and volume percentage (v/v%). Each melting temperature *t*_m_ is obtained by thermal unfolding of the proteins at a given concentration of FA or HCl followed by fitting the sigmoidal unfolding curves eq. (1) (Fig. 2). *t*_m_ values for all three proteins fall significantly at high FA concentrations compared to HCl solutions of the corresponding pH.

**Figure 4.**
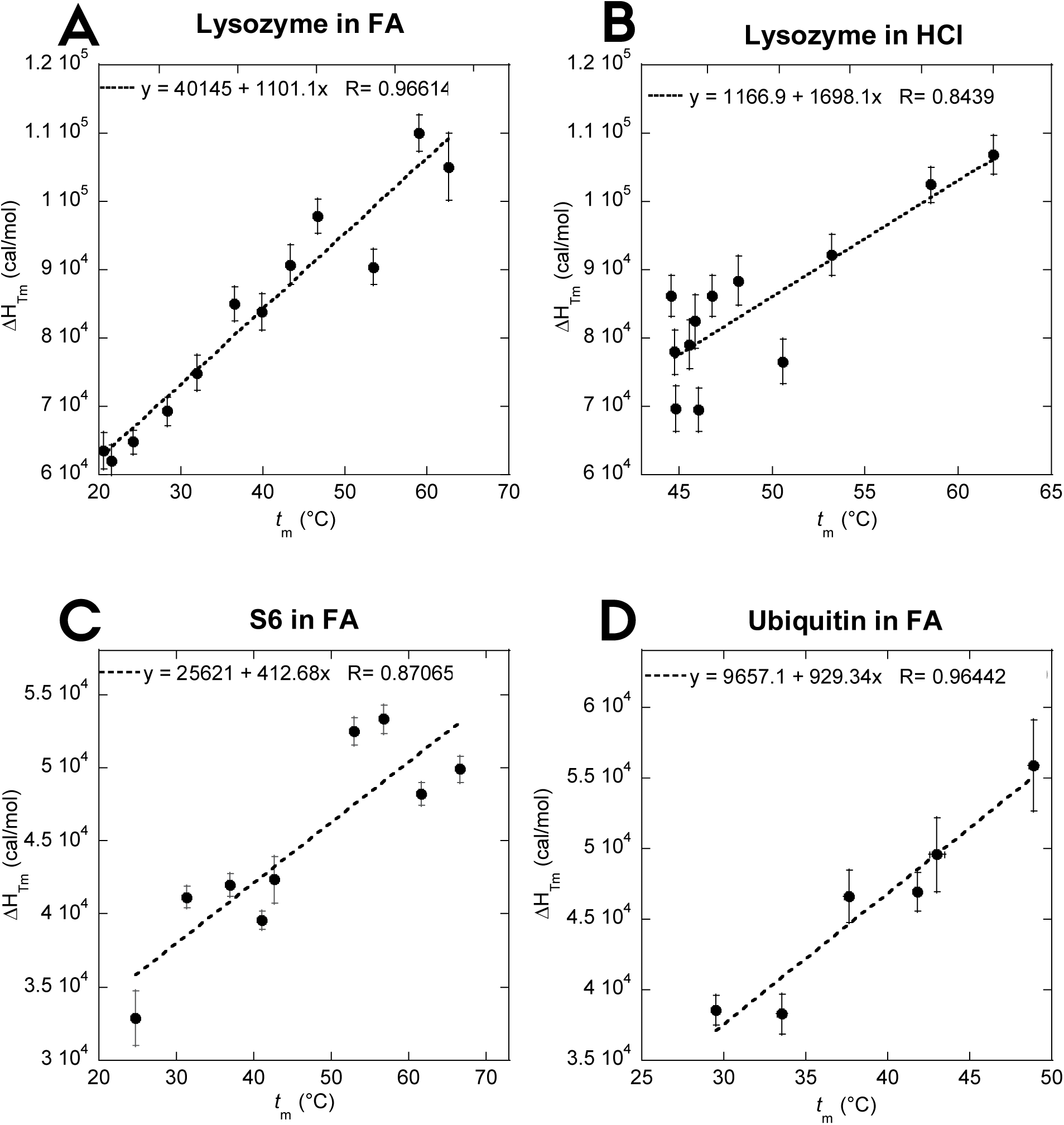
Determination of the specific heat capacity of denaturation (ΔC_p_) of lysozyme in (A) FA and (B) HCl, (C) S6 in FA and (D) ubiquitin in FA by plotting Δ*H*_Tm_ versus *t*_m_. ΔC_p_ of S6 and ubiquitin in HCl could not be determined due to insignificant difference in measured t_m_-values at different concentrations of HCl.

### Heat capacity changes upon unfolding are reduced in FA compared to HCl and GdmCl, which may reflect a more compact unfolded state and a more expanded native state

To quantify this denaturation effect, we calculated the free energy of unfolding (Δ*G*_D-N_) of the different proteins in HCl and FA at matching pH values and subtracted the effect of HCl. To achieve this, we needed to calculate the specific heat capacity change upon unfolding (ΔC_p_) of the different proteins under our denaturation conditions. Standard thermodynamics predicts a linear relationship between *T*_m_ and ΔH_*T*m_ with a slope of ΔC_p_ (assuming that ΔC_p_ is temperature-independent)^*54*^. Indeed, plots of ΔH_*T*m_ versus *t*_m_ provide reasonably linear plots (Fig. 4). Based on these plots, ΔC_p_ of HEWL is calculated to 1.10 ± 0.10 kcal mol^-1^K^-1^ in FA and 1.70 ± 0.34 kcal mol^-1^ K^-1^ in HCl. An early calorimetric study of HEWL denaturation reported a ΔC_p_ of 1.37 kcal mol^-1^ K^-1^ when the protein is denatured in guanidinium chloride (GdmCl), but a similar ΔC_p_ of 1.57 kcal mol^-1^ K^-1^ when denatured in in HCl^*55*^. FA is reported to induce a more compact state when used as a solvent^*30*^ and considering that the ΔC_p_ value is correlated to the change in solvent-accessible surface area (SASA) upon unfolding^*56*^ (*i.e.* the change in compaction upon unfolding) this may explain the difference in ΔC_p_ values between FA and HCl.

For both S6 and Ubi, we determined ΔC_p_ using FA-based scans, since the small variation in *t*_m_ with pH in the HCl-based thermal scans prevented the determination of meaningful *t*_m_ - ΔH_*T*m_ plots. The ΔC_p_ of S6 in FA is found to be 0.41 kcal mol^-1^K^-1^, which is significantly lower than the value of 0.93 ± 0.05 kcal mol^-1^K^-1^ obtained when denaturing with GdmCl^*57*^. We attribute this to the previously mentioned “loosening” of the native state of S6 at low pH, which – together with a likely more compact unfolded state in FA compared to GdmCl – means that the increase in SASA upon unfolding will be smaller. Nevertheless, the satisfactory linear correlation between *t*_m_ and ΔH_*T*m_ for S6 is consistent with a well-defined reasonably compact species populated prior to thermal unfolding.

Somewhat surprisingly in view of the previous results, ΔC_p_ of Ubi in FA was found to be 0.93 ± 0.13 kcal mol^- 1^K^-1^, which is essentially identical to the value of 0.90 ± 0.20 kcal mol^-1^K^-1^ obtained from calorimetric studies in different concentrations of GdmCl^*48*^.

### pH effects can be factored out to reveal the intrinsic contribution of formic acid to protein stability, which is 3-fold weaker than GdmCl

Having determined ΔC_p_, Δ*G*_D-N_ can be calculated from each thermal scan based on their individual ΔH_*T*m_ and *T*_m_ values (eq. 2). For all three proteins, Δ*G*_D-N_ (here denoted Δ*G*D-N^FA+pH^ to emphasize the joint contribution of FA as a chemical denaturant and as an acid) decreases linearly with [FA] (blue points and lines in Fig. 5). Each FA-Δ*G*_D-N_ data point has a HCl-Δ*G*_D-N_ counterpoint, in which Δ*G*_D-N_ has been determined from a thermal scan performed in a HCl solution with the same pH-value as the corresponding FA data point (Δ*G*D-N^pH^, red points and lines in Fig. 5). The low Δ*G*_D-N_-values of the HCl-based data points compared to the FA-based points make it clear that the stability decrease caused by a given concentration of FA (which can be considered to be driven by the combined effect of lowered pH and increased [FA]) is significantly larger than the stability decrease solely caused by the equivalent decrease in pH. This emphasizes the strong effect of FA as a denaturant, and not merely as an acid. Thanks to these pairwise measurements of stability in FA and HCl at the same pH-value, it is possible to subtract the isolated pH-effect for FA at every data point and thus calculate the “true” FA effect, denoted Δ*G*D-N^FA^, as a function of [FA]. This corrected series is shown in green in the graphs in Fig. 5. For all three proteins, there is a satisfactory linear relationship between Δ*G* D-N^FA^ and [FA], whose slope is the *m*_FA_ value. Lysozyme’s *m*_FA_-value of 1.28 ± 0.10 kcal mol^-1^M^-1^ is close to the *m*-value of 1.29 kcal mol^-1^M^-1^ reported for lysozyme when denatured in urea^*56, 58*^, but significantly lower than the value of 3.96 ± 0.47 kcal mol^-1^M^-1^ in GdmCl^*59*^. A similar analysis for S6 yields an *m*_FA_-value of 0.81 ± 0.12 kcal mol^-1^M^-1^ which is again considerably smaller than the value of 2.33 ± 0.12 kcal mol^-1^M^-1^ in GdmCl^*60*^ (note that this value is based on the average *m*-value of 1.71 ± 0.09 M^-1^ from ^*60*^ and has to be multiplied by –*RT* ln10). Finally, the *m*_FA_-value of Ubi in FA was found to 0.59 ± 0.11 kcal mol^-1^M^-1^, which compares to 1.35 kcal mol^- 1^M^-1^ in GdmCl^*44*^. *m*-values reflect the molar efficacy of a given denaturant in reducing the stability of a protein, and can be correlated to effects such as the ability to increase solubility of nonpolar amino acids, that are normally buried within a protein core, as well as hydrogen bonding to peptide bonds and other polar groups in proteins^*61-63*^. Such values may be further substantiated by *e.g.* measuring the free energy of transfer of individual amino acids and small peptides to the denaturant in question to determine solubility effects^*64*^. Overall the *m*_FA_ values are ∼ 1/3 of the corresponding *m*_GdmCl_ values, illustrating that FA is only 1/3 as efficient as GdmCl as a denaturant on a per-mole basis (discounting the pH effect).

**Figure 5.**
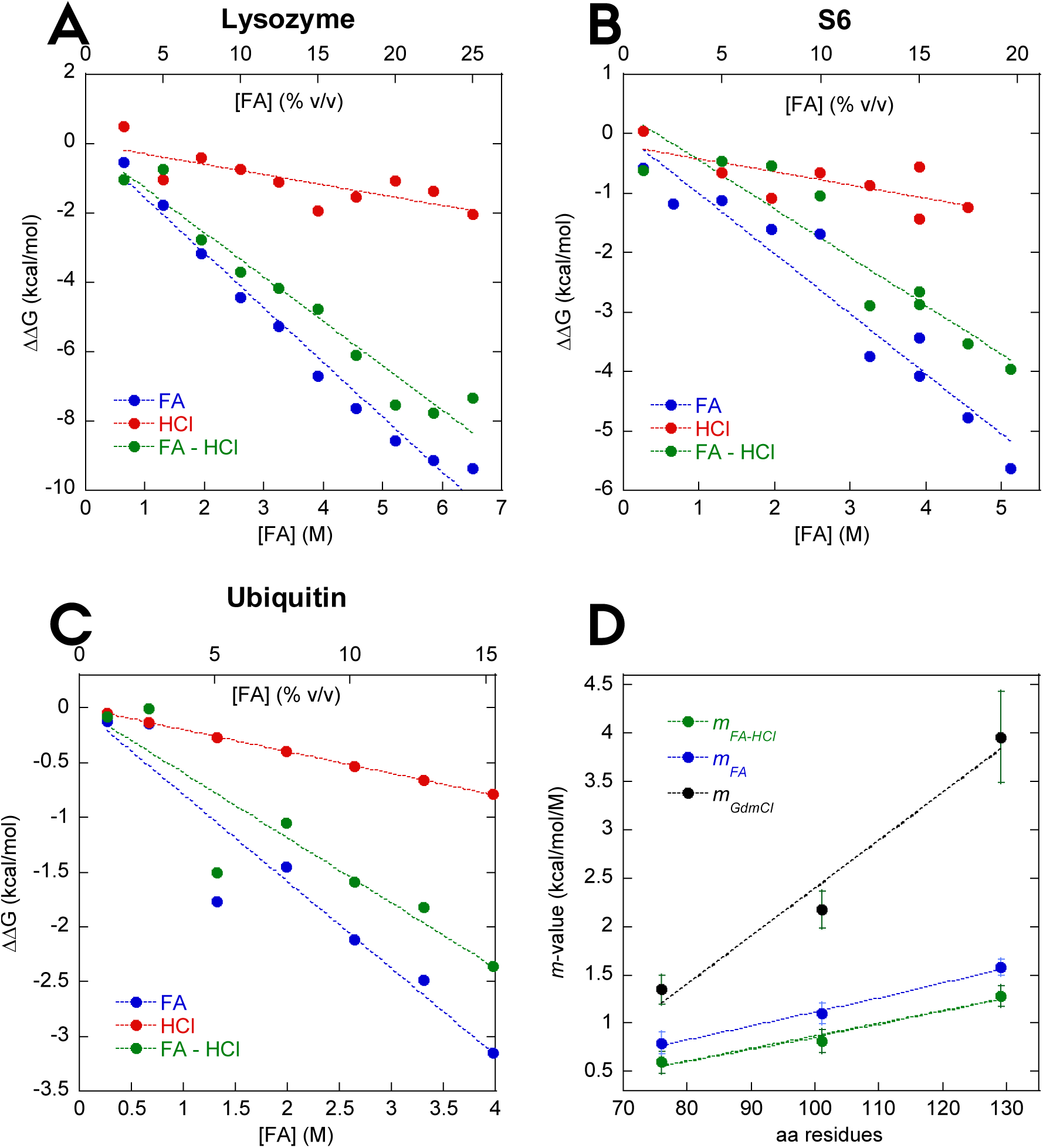
Correlation between the change in the free energy of unflding ΔΔG_D-N_ and FA concentration (blue), HCl (red) and HCl-subtracted effect of FA (green) of (A) lysozyme, (B) S6 and (C) ubiquitin. (D) The *m*-values found in these panels are plotted versus the number of residues of the model proteins. The *m*-values scale with the protein size of the three proteins analogously to GdmCl, where its slope is 2.6 times more steep, reflecting FA’s comparatively lower strength as a denaturant.

### Isothermal titration with formic acid confirms results from thermal scans

As a final control, we sought to validate the obtained *m*-values above through an analysis of FA-induced denaturation under isothermal denaturation (25°C) measured by intrinsic aromatic fluorescence. Note that we observe only *m*_FA+pH_ values, and not the corrected *m*_FA_, as the pH-effect cannot be subtracted in a simple manner in this type of experiment. In addition, we also measure [FA]^50%^, which is the FA concentration at which half of the protein is unfolded. We were able to predict the fraction of folded protein in FA at isothermal conditions based on the Δ*G*_D-N_^FA+pH^ and *m*_FA+pH_ values from the thermal unfolding experiments using eq. (3) (Fig. 6A-C). For all 3 proteins, there is excellent overlap between isothermal data and folding fractions predicted from thermal scans. For both lysozyme and Ubi there is a good agreement between *m*_FA+pH_ obtained from both thermal and isothermal experiments. Lysozyme unfolding, represented by a red-shift in Trp fluorescence, overlaps the predicted *f*_N_ very well and shows an *m*_FA+pH_ value of 1.43 ± 12 kcal mol^-1^ M^-1^, which is close to the value of 1.58 ± 0.08 kcal mol^-1^ M^-1^ measured by near-UV CD (Fig. 6A). Ubi unfolding was followed as a change in emission at 320 nm (Trp+Tyr) and overlaps the predicted curve almost perfectly (Fig. 6C). Both thermal and isothermal measurements of Ubi unfolding show a strikingly similar *m*_FA+pH_ value of 0.79 ± 0.11 kcal mol^-1^ M^-1^. Isothermal measurements of S6 was measured by the change in near-UV ellipticity to avoid complications from protonation of Trp that affect fluorescence without affecting structure. The isothermal *m*_FA+pH_ value of 0.66 ± 0.22 kcal mol^-1^ M^-1^ is within error the same as the value of 0.81 ± 0.12 kcal mol^-1^ M^-^1 from thermal unfolding (Fig. 6B). These values are summarized in Table 1.

**Table 1.** Thermodynamic parameters of lysozyme, S6 and ubiquitin denaturation in FA derived from the thermal and isothermal unfolding experiments measured by near-UV circular dichroism and aromatic fluorescence, respectively.

| Method | Parameter | Lysozyme | S6 | Ubiquitin |
| --- | --- | --- | --- | --- |
| **Near-UV CD**<br>**(thermal)** | $\Delta C_p^{FA}$ (cal mol<sup>-1</sup> K<sup>-1</sup>) | 1101 ± 93 | 412.7 ± 88 | 928.3 ± 13 |
| | $m_{FA+pH}$ (kcal mol<sup>-1</sup> M<sup>-1</sup>) | 1.58 ± 0.08 | 1.10 ± 0.11 | 0.79 ± 0.11 |
| | $m_{FA}$ (kcal mol<sup>-1</sup> M<sup>-1</sup>) | 1.28 ± 0.10 | 0.81 ± 0.12 | 0.59 ± 0.11 |
| **Fluorescence**<br>**(isothermal)** | [FA]<sup>50%</sup> (M) | 5.18 ± 0.05 | 4.86 ± 1.3 | 4.69 ± 0.40 |
| | $m_{FA+pH}$ (kcal mol<sup>-1</sup> M<sup>-1</sup>) | 1.43 ± 12 | 0.66 ± 0.21 | 0.79 ± 0.11 |

**Figure 6.**
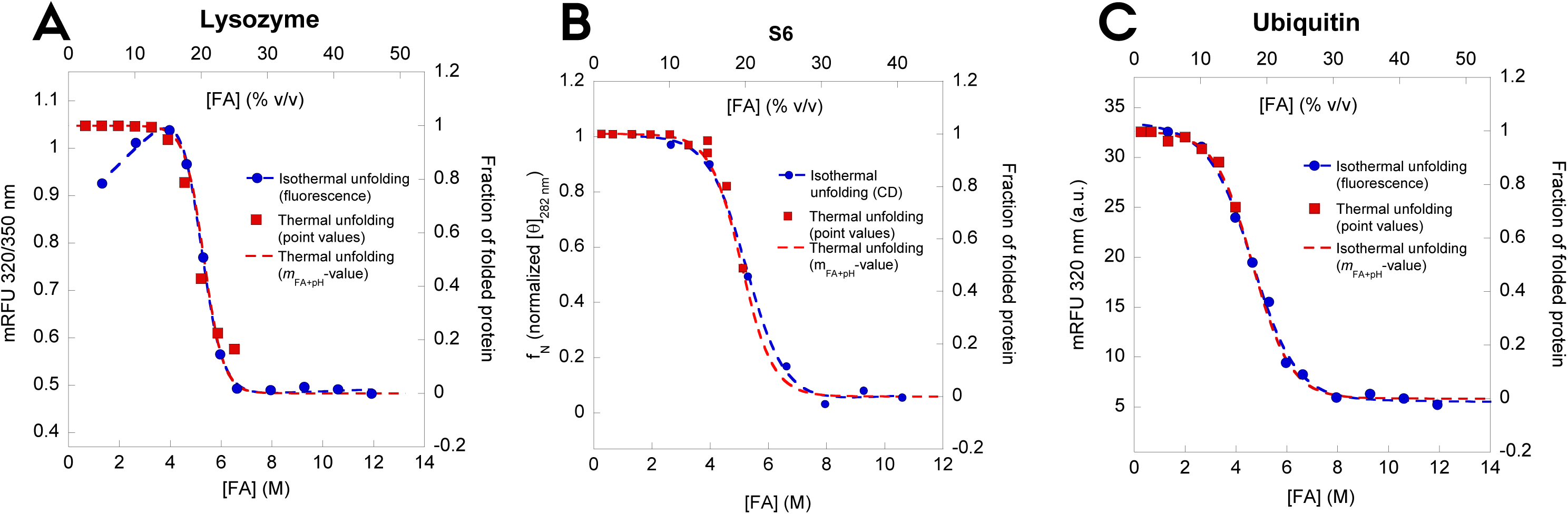
Unfolding of (A) lysozyme, (B) S6 and (C) ubiquitin measured by intrinsic aromatic fluorescence (lysozyme and ubiquitin) and near-UV CD (S6). Blue points are measured equilibrium values while red points represent the predicted fraction of folded protein at 25°C based on thermal unfolding values. The red stippled curves are best fits of eq. 4 to the red points. These show a very good correspondence with the measured isothermal equilibrium dataset.

Gratifyingly, the *m*-values of all three proteins scale with their size measured as residue number (Fig. 5D). Similarly, *m*-values for denaturation with classical denaturants such as urea and GdmCl scale linearly with the change in SASA upon unfolding, which in turn scales with the number of residues per protein^*56*^, as shown in Fig. 2D. These linear correlations are attributed to weak binding effects, such that the interaction between the denaturant and the protein is determined by the SASA rather than specific interaction regions. This can also be quantified as the increase in solubilisation power of the denaturant, *i.e.* their interactions with protein functional groups which become exposed upon unfolding. Thus, the better the solvation, the stronger the denaturation potency, cfr. the seminal work by Nozaki and Tanford on the solubility of amino acids in GdmCl^*64*^ which has subsequently been used to develop activity coefficients for GdmCl denaturation^*59*^. FA must work in the same unspecific manner given its scaling properties, indicating that it has a conventional denaturing effect as weak binder to the protein surface in addition to the effect of lowering the solution pH. However, the slope of the plot for GdmCl (−0.049 ± 0.009 kcal/mol/M/residue) is 3-4 times steeper than that of FA (0.013 ± 0.002 kcal/mol/M/residue), illustrating the greater solubilisation power of GdmCl (Fig. 5D).

### Formate does not show any denaturating potency and formamide is ∼3-fold weaker as denaturant than formic acid, indicating lower solvating abilities

We also attempted to quantify the denaturation potency of the ionized version of FA, sodium formate, as well as its amidated counterpart formamide (FM). However, thermal denaturation of HEWL in formate was challenged by the high levels of precipitation accompanying unfolding (Fig. S3AB). In addition, the *t*_m_ of unfolding increased from 72.4°C (0 M formate) to 78-80°C (0.8-2.6 M formate) (Fig. S3C). This stabilization by formate indicates that only the protonated form of FA has denaturing properties. FM, on the other hand, showed a more classical denaturation effect (Fig. S4A) and in addition, denaturation was completely reversible up to 45%, above which it became progressively less reversible (data not shown). *t*_m_ decreased linearly with [FM] up to 80% (Fig. S4B). A plot of ΔH_*T*m_ as a function of *t*_m_ was satisfactorily linear (Fig. S4C), leading to a predicted HEWL ΔC_p_ of 1.25 ± 0.11 kcal mol^-1^K^-1^, which is identical within error to the value of 1.10 ± 0.10 kcal mol^-1^K^-1^ in FA, suggesting a comparable level of unfolding and thus hydration in the denatured state. In the FM concentration range we probed here (0-80%, corresponding to 0-23 M) the pH of the solution only increased from 8.45 to 10.1 and, unsurprisingly, the *T*_m_ of lysozyme only decreased by ∼ 6°C in the corresponding control experiments where pH was adjusted with NaOH buffer. The very modest decline in stability induced by the shift in pH alone was nevertheless subtracted from the overall Δ*G*_D-N_, analogous to Fig. 5A-C. This led to a linear correlation between Δ*G*_D-N_ and [FM] (Fig. S4D) with an *m*_FM_-value of 0.47 ± 0.02 kcal mol^-1^M^-1^ which is 2.7-fold less than *m*_FA_ for lysozyme (1.28 ± 0.10 kcal mol^-1^M^-1^). Equilibrium fluorescence measurements at isothermal conditions show an even lower estimate of *m*_FM_-value of 0.29 ± 0.06 kcal mol^-1^M^-1^ (Fig. S4E), though the denaturation curve and the curve constructed from thermal scans overall overlap well. Thus, FM is significantly less efficient than FA as a denaturant even though the only difference between these two molecules is the replacement of a carboxylic acid with an amide group. Therefore we conclude that the acidic -OH group of FA must interact with and solvate protein functional groups (particularly amides and hydrocarbon groups) better than the amide -NH_2_ group (cfr. similar solubilisation measurements for amide-amide and amide-hydrocarbon interactions)^*65*^. Similar considerations apply for –OH (FA) versus – O^-^ (formate). Future studies using appropriate model compounds may put these empirical observations on an even firmer basis.

### Solubilization of FapC variants by formic acid reveals that FapC imperfect repeats stabilize fibrils

Having quantified the denaturating potency of FA, we now analyse the contribution of the different imperfect repeats to FapC fibril stability. We designed different FapC mutant proteins where either one repeat (FapC ΔR1, FapC ΔR2 and FapC ΔR3), two repeats (FapC ΔR1R2, FapC ΔR1R3 and FapC ΔR2R3) or all three repeats (FapC ΔR1R2R3) were removed (Fig. S1). These proteins were fibrillated and their secondary structure was analysed with FTIR. Consistent with previous observations^*24, 66*^, all fibrils showed an intense peak around 1620 cm^-1^, indicating that they contain the characteristic amyloid cross-β structure as opposed to conventional β-sheets which absorb at wavenumbers > 1630 cm^-1^ (Fig. 7A). This is caused by the stacking of the β-sheets in the fibril structure which shifts the frequencies towards lower wavenumbers^*67*^.

**Figure 7.**
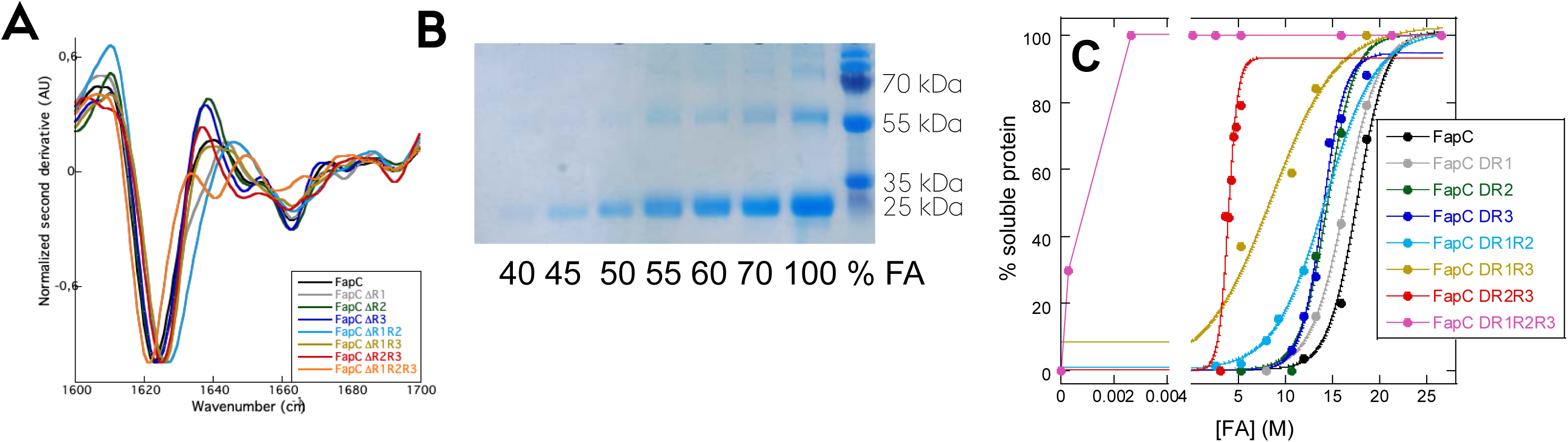
All FapC proteins can form amyloid fibrils. (A) FapC variants were desalted from 8 M GdnCl into milliQ water and immediately fibrillated before FTIR analysis was performed on the formed fibrils. The normalized second derivative is shown. (B) Representative gel of the supernatants after treatment with FA (here shown for FapC ΔR3 protein). The band around 25 kDa is FapC solubilized by FA. (C) Percentage total protein present in supernatant after FA treatment as a function of FA concentration. Gel bands were quantified with ImageJ, normalized to the 100% FA band and fitted to eq. 6. There is a marked decrease in stability when the number of repeats is reduced.

We used FA to measure the stability of the different FapC constructs as follows. Equal amounts of fibril mass were treated with different concentrations of FA and the amount of fibrils that were dissolved by FA was quantified from SDS-PAGE gels (an example is shown in Fig. 7B). All quantifications were normalized to the sample treated with 100% FA where the fibrils are fully dissolved. FA treatment of all eight proteins showed a stepwise displacement of the midpoint values of denaturation ([FA]^50%^) towards lower values of FA as more and more repeats were removed (Fig. 7C, summarized in Table 2). The stability towards FA is dependent not only on *how many* repeats but also on *which* repeats are removed from the protein. Based on the decline in [FA]^50%^, removal of R1 has a relatively modest impact on fibril stability compared to R2 and R3. In good agreement with these results, bioinformatic investigations of the sequences for the three repeats using the FISH Amyloid tool^*68*^ show that R1 is the least aggregation-prone repeat (and by implication contributes the least to fibril stability), with an average amyloidogenicity score pr. amino acid of 0.0353 compared to 0.0588 for R2 and 0.0437 for R3 (Fig. S5). However, its *m*-value is slightly decreased compared to the two other mutants, leading to a lower overall stability. Importantly, removing two repeats leads to a significant decrease in either [FA]^50%^ (ΔR1R3 or ΔR2R3) or the *m*_FA_-value (ΔR1R2 and ΔR1R3) and a consequent dip in the overall stability of the protein. Besides the FapC ΔR1R2R3 mutant (see below), fibrils formed from the FapC ΔR2R3 and FapC ΔR1R3 proteins are the most destabilized compared to the wt FapC, indicating an important role for R3 in stabilizing the fibrils. That the different FapC repeats play different roles in fibril formation has also been observed for CsgA where the last repeat (R5) is the most aggregation-prone and also very critical for polymerization^*22*^.

**Table 2.**
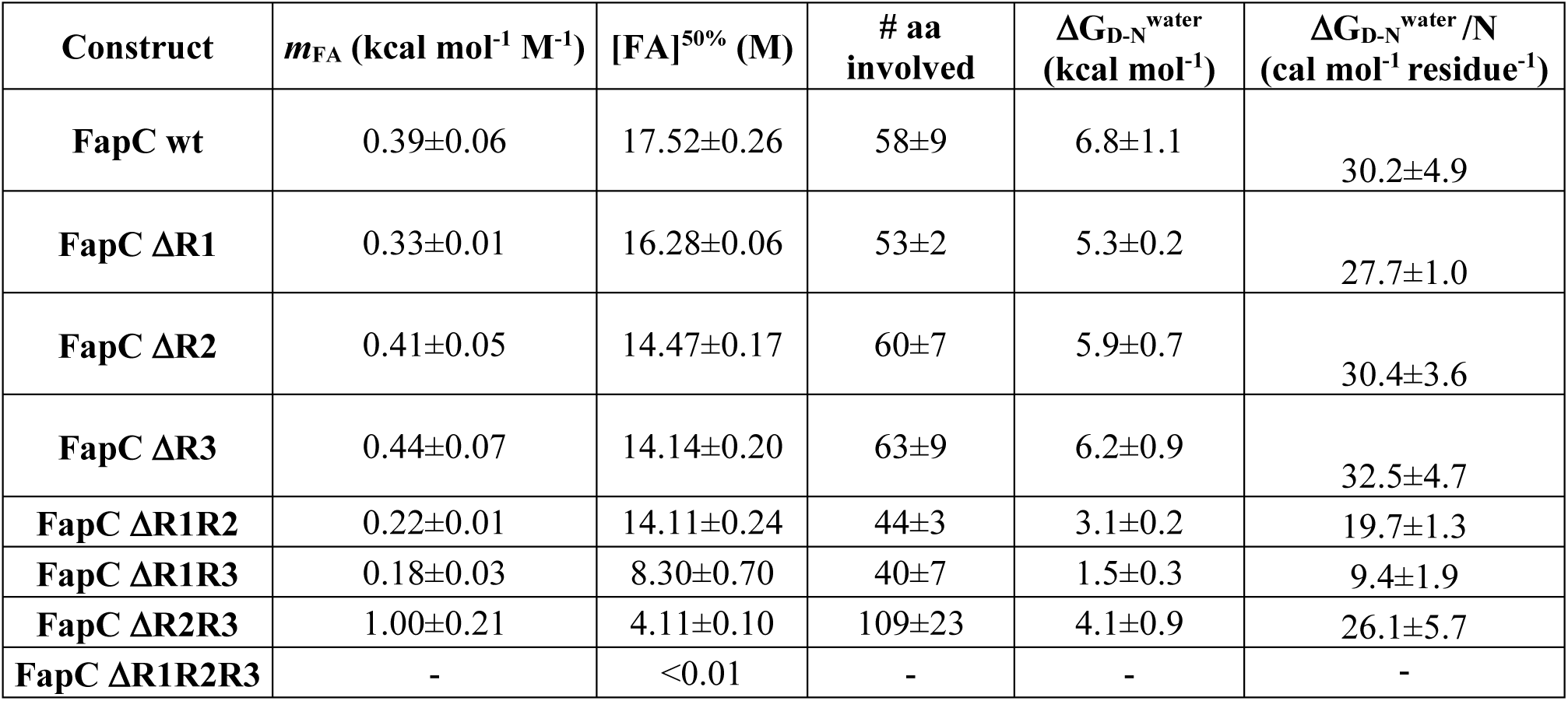
Summary of FA stability of different FapC constructs based on solubilization measurements.

| Construct | $m_{FA}$ (kcal mol<sup>-1</sup> M<sup>-1</sup>) | [FA]<sup>50%</sup> (M) | # aa involved | $\Delta G_{D-N}^{water}$ (kcal mol<sup>-1</sup>) | $\Delta G_{D-N}^{water}/N$ (cal mol<sup>-1</sup> residue<sup>-1</sup>) |
| --- | --- | --- | --- | --- | --- |
| **FapC wt** | 0.39±0.06 | 17.52±0.26 | 58±9 | 6.8±1.1 | 30.2±4.9 |
| **FapC ΔR1** | 0.33±0.01 | 16.28±0.06 | 53±2 | 5.3±0.2 | 27.7±1.0 |
| **FapC ΔR2** | 0.41±0.05 | 14.47±0.17 | 60±7 | 5.9±0.7 | 30.4±3.6 |
| **FapC ΔR3** | 0.44±0.07 | 14.14±0.20 | 63±9 | 6.2±0.9 | 32.5±4.7 |
| **FapC ΔR1R2** | 0.22±0.01 | 14.11±0.24 | 44±3 | 3.1±0.2 | 19.7±1.3 |
| **FapC ΔR1R3** | 0.18±0.03 | 8.30±0.70 | 40±7 | 1.5±0.3 | 9.4±1.9 |
| **FapC ΔR2R3** | 1.00±0.21 | 4.11±0.10 | 109±23 | 4.1±0.9 | 26.1±5.7 |
| **FapC ΔR1R2R3** | - | <0.01 | - | - | - |

The repeat-less mutant (FapC ΔR1R2R3) is extremely sensitive towards FA; already at 0.001% FA (0.3 mM) 30% is dissolved, and at 0.1% FA there is no amyloid left. This mutant is composed entirely of FapC’s linker regions but nevertheless still forms fibrils on the hours-days timescale, although the fibrillation process is much less reproducible than for the other FapC constructs^*24, 66*^. Clearly the fibrils formed by this mutant are much less resistant towards FA than any of the other mutants, in fact its properties are similar to fibrils formed by misfolded proteins such as α-synuclein, which dissolves completely around 0.1% FA (data not shown). Thus fibrils containing as little as one single repeat retain a moderate resistance to FA, while there is a dramatic collapse of stability upon complete removal of repeats. This strongly indicates that repeats are indeed the key to functional amyloid stability.

Analysis of the imperfect repeat sequences using the ZipperDB tool^*69*^ reveals that both the N-terminal part (excluding the signal peptide), both linkers (L1 and L2) and the C-terminal have segments of high fibrillation propensity (below −23 kcal/mol) (Fig. S6). These segments may rationalize the ability of this highly amputated mutant to form fibrils with amyloid characteristics, although the ensuing fibrils’ high sensitivity to FA indicates that the linkers only make minor contributions to the exceptional stability of the functional amyloid. Indeed, linker regions are completely missing from the functional amyloid in *E. coli*, CsgA, whose five repeats are connected by only enough residues (4-5) to form a tight turn^*22*^.

### Solubility in SDS correlates well with stability against FA

To obtain complementary evidence for the quantification of destabilization of the different FapC variants, we turned to solubilisation in SDS. FapC fibrils are largely resistant towards strongly denaturing anionic surfactants such as SDS; boiling SDS is even used to remove less robust proteins as a way to purify FapC fibrils^*13*^. Nevertheless, the effect has not been investigated systematically and even modest solubilisation levels will be indicative of fibril stability. Accordingly, we incubated FapC fibrils in 58 mM SDS (corresponding to the concentration of SDS in SDS-PAGE loading buffer when loading samples on a gel) at a range of temperatures (21-95°C) for 10 min. Supernatants from these incubations were then analysed by SDS-PAGE to determine the amount of soluble protein. We observed a strong temperature dependence, with estimated solubilities rising from ∼ 0.2% at 21°C to ∼ 5% at 95°C (Fig. 8A). Solubility was only slightly increased under reducing conditions (1 mM TCEP, Fig. 8A). FapC contains two conserved Cys residues in a CXXC motif near the C-terminus (position 213 and 216, Fig. S1) which may form intermolecular disulfide bonds during the early on-pathway FapC oligomerization steps, but this does not impact stability significantly^*70*^. We carried out similar studies with the other 7 FapC deletion mutants and observed a significant increase in solubility for several of the mutants (Fig. 8B), up to 30-35% for the double mutant FapC ΔR2R3 and the triple mutant FapC ΔR1R2R3. We observe a very significant correlation between solubility in SDS at 20°C (both under reducing and oxidizing conditions) and the midpoint of denaturation in FA (Fig. 8C). This corroboration by an independent approach indicates that solubility in FA is indeed a credible way of estimating fibrils’ intrinsic ability to dissociate.

**Figure 8.**
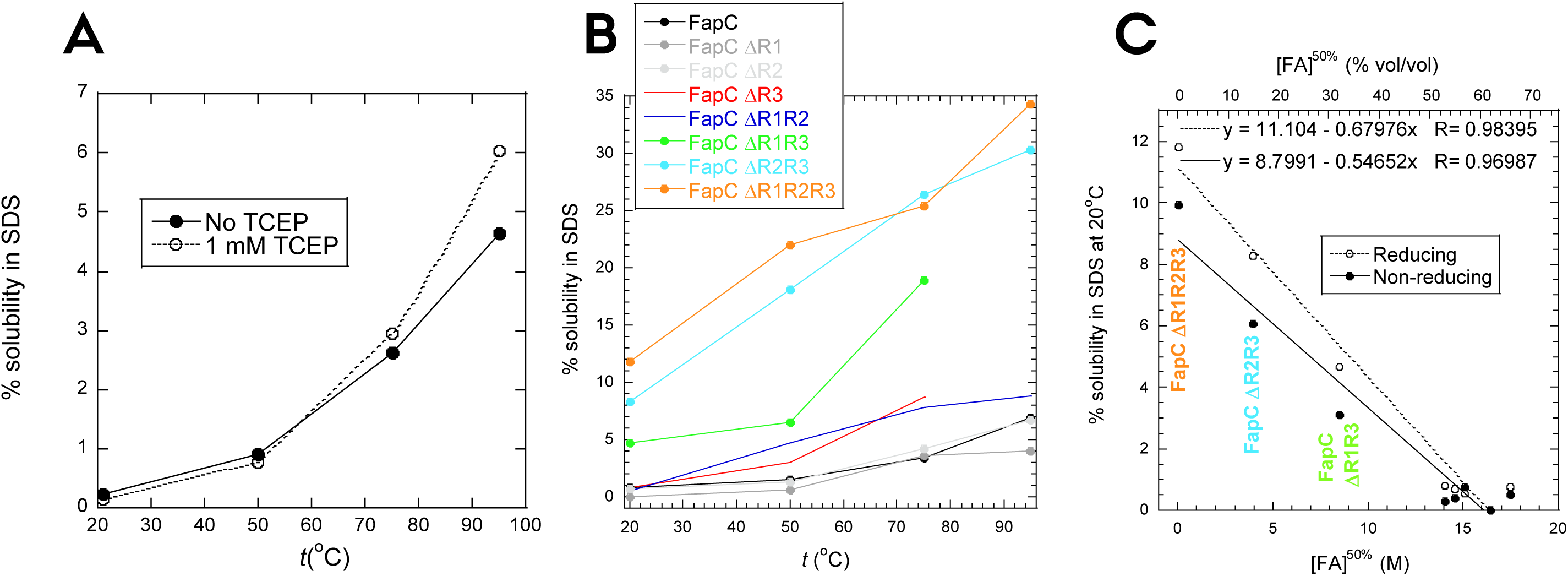
Solubilization of FapC fibrils in SDS. (A) Solubility of wt FapC in SDS at 20-95°C under reducing (1 mM TCEP) or non-reducing conditions. Solubility was quantified as in Fig. 7C. (B) Solubility under reducing conditions in SDS at 20-95°C for all 8 FapC variants. (C) Correlation between solubility in SDS at room temperature and the midpoint of denaturation in FA, shown for all 8 FapC variants. The three most destabilized variants are indicated with the same colors as in panel B.

### FapC fibril solubilization by FA predicts a relatively low stability compared to other fibrils; superior performance by FA is not straightforward to explain

We transformed the solubility data of FapC mutants in Fig. 7 to apparent stability as follows. Based on our previous analysis of the stabilities of HEWL, S6 and Ubi in FA, we assume a linear relationship between [FA] and the free energy of unfolding, ΔG_D-N_. The associated equilibrium constant of unfolding is formally defined as the ratio between dissolved and aggregated protein. We also considered the alternative option of assuming that the equilibrium constant corresponds to the concentration of free protein, inspired by the use of free monomer concentration to determine the stability of Aβ fibrils^*25*^ as well as other fibril stability studies^*26*^; however, this approach does not yield a linear relationship between the free energy and [FA] and therefore does not allow us to extrapolate robustly to 0 M FA (data not shown). Data are instead fitted with eq. 2 and summarized in Table 2. In line with the general decline in values of [FA]^50%^, they reveal a remarkable reduction in stability upon removal of two repeats. The decrease is less significant when only one repeat is removed, due to the errors associated with the measurement, but they still reveal a reducing trend, particularly when viewed from the perspective of the midpoint of denaturation.

The overall stability of the fibrils (6.8 kcal/mol for full-length FapC and less for the variants) is very modest compared to naturally occurring globular proteins which have stabilities in the 5-15 kcal/mol range^*71*^. The value is comparable to, but certainly not higher than, the stabilities of fibrils formed from a range of other proteins which are associated with neurodegenerative diseases or other misfolding conditions^*26*^ but not otherwise optimized to form fibrils. The free energy per residue for FapC (ca. 30 cal/mol/residue) is actually substantially lower than any of the measured stabilities for these proteins and peptides, for which the values vary from ca. 40 to 500 cal/mol/residue. The highest stability contribution per residue in this study is assumed by small peptides whose entire sequence is most likely integrated into the amyloid structure, while larger proteins with more extensive regions of non-amyloid structure lead to lower scores^*26*^. For FapC, it is likely that the linker regions are not involved in amyloid formation^*72*^; however, restricting the analysis to the 3 imperfect repeats as contributors to fibril stability only doubles this figure and still leaves it in the low end of the scale. Assuming that our method of fibril stability estimation is correct, this raises the question why a chemical denaturant such as GdmCl, which effectively unfolds very stable proteins such as OmpA (ΔG_D-N_ = 16 kcal/mol and t_½_^unfolding^ at 0M GdmCl ∼ 800 years^*73*^), is quite unable to dissolve these fibrils. The point is made all the stronger by the fact that *m*-values for GdmCl based on our 3 globular proteins are ∼ 3 times higher than for FA, illustrating the high efficacy of solubilization by GdmCl. Nevertheless, even a crass anionic surfactant like SDS only confers limited solubilization at temperatures as high as 95°C. We speculate that the unique solubilization properties of FA may reflect (1) a high activation barrier for FapC unfolding which is selectively reduced by FA due to favorable solvation of the transition state, (2) very favorable solvation of the denatured state or (3) a combination of the two phenomena. In the absence of kinetic data (which are currently not feasible to record *e.g.* by near-UV CD due to the high concentrations of FA needed and the low near-UV CD signal of FapC fibrils), it is not possible to provide any direct support for transition state stabilization, but it is worth noting the cooperative nature of the fibril build-up which in itself constitutes a significant activation barrier to dissociation. Thus, each FapC monomer is stabilized not only by interactions with other monomers in the fibril (at least two monomers on either side in the fibril and possibly also inter-sheet contacts between multiple protofilaments in a given fibril) but also by contacts between the individual imperfect repeats which most likely constitute β-hairpin helices^*21, 23*^.

### Using formic acid stability to predict the level of involvement of amino acids in fibrillation

Based on the correlation in Fig. 5D between *m*_FA_ and protein size, we use the *m*_FA_-values in Table 2 to predict the number of residues involved in FapC amyloid formation (Table 2). This assumes that the average degree of burial per residue (*i.e.* the reduction in solvent accessible surface area) is the same for the 3 globular proteins as for the residues forming amyloid structures. Such an assumption may be questioned. Loops can remain significantly exposed in the native state, and generally β-sheet proteins bury more surface area than α-helices and turns/loops, leading to ∼10, 20 and 30% accessibility, respectively, compared to the denatured state^*74*^. A higher level of burial per residue means that a smaller number of residues is required to achieve the same level of burial, and therefore transfer of data based on *e.g.* pure α-helical properties to analysis of β-sheet proteins may lead to an overestimate of the number of residues involved in a given folding process. Both lysozyme, S6 and ubiquitin contain significant amounts (10/46/34%, respectively) of β-sheet structure but also a major proportion of less organized structure like coils and turns (49/27/43%, respectively), leading to a lower level of burial than all-β-sheet structures.

Full length FapC and all single-repeat mutants show the same overall level of amino acids involved (∼60); this declines to ∼40 for two double repeats but then rises steeply to 110 for the third double repeat (FapC ΔR2R3). The number of residues in the imperfect repeats are ca. 105 for full-length FapC, declining to 70 for single repeats and 35 for double repeats. Thus there is an overall satisfactory prediction for single repeats and two double repeats (with the caveats on *m*_FA_-values described above). The breakdown in the relationship between repeat length and *m*-values for FapC ΔR2R3 may be caused by a change in the nature of the fibril structure relative to the FA-denatured state. A greater *m*-value indicates a greater difference between compactness of the two ground states (FA-denatured and fibrillar states). Thus, either or both of the two states could change compactness; the denatured state could become more expanded and/or the fibrillar state more compact. The denatured state of FapC in FA has not been characterized in any detail, but we note that aggregated species are found at the very beginning of the aggregation of wt FapC according to Small Angle X-ray Scattering^*75, 76*^. This might be explained by a high propensity even in the denatured state to engage in intermolecular contacts; removal of repeats could destabilize these transient interactions and lead to a more fully expanded state. It seems unlikely that removal of fibrillar repeats would lead to a more compact structure unless the protein changes architecture to a fundamental extent, *e.g.* by integrating otherwise exposed regions such as the linker sequences into the fibrillar state. This could be a possibility in view of the ability of the repeat-free mutant to form (admittedly very FA-sensitive) fibrils^*24, 77*^. More detailed but also methodologically demanding studies on the denatured and fibrillated states of FapC ΔR2R3, as well as the kinetics of their interconversion, are needed to address these questions.

## CONCLUSION

We have quantitated the denaturing potency of formic acid against globular proteins using thermal scans and isothermal titration monitored by near-UV CD. FA unfolds globular proteins reversibly and cooperatively; the effects go beyond simple pH-driven denaturation and show a linear relationship between unfolding free energy and FA concentration, revealing that FA is ∼3-fold less effective on a molar scale than guanidinium chloride as a denaturing agent. Nevertheless, protonated FA is radically different from its ionized formate counterpart, which actually stabilizes proteins. Formamide is ca. 3 fold weaker than FA as denaturant, highlighting the role of the acidic -OH group. Heat capacity changes indicate that the unfolded state is more compact in FA than in chemical denaturants. The *m*_FA_-value of unfolding scales with protein size, suggesting weak binding interactions between FA and the polypeptide chain. FA was used to solubilize fibrils of the functional amyloid FapC, revealing a surprisingly low stability. Importantly, stepwise removal of the imperfect repeats progressively decreased stability. Complete removal, while not preventing fibrillation, led to fibrils that were extraordinarily sensitive to FA, indicating that these repeats are central to the stability of functional amyloid.

## EXPERIMENTAL PROCEDURES

### Materials

Hen egg white lysozyme (HEWL), bovine ubiquitin (Ubi) and all other chemicals were from Sigma-Aldrich.

### Design and purification of the FapC repeat deletion mutant proteins

Seven FapC mutant proteins (based on FapC from the *Pseudomonas sp.* UK4 strain) lacking 1-3 of the imperfect repeats (FapC ΔR1, ΔR2, ΔR3, ΔR1R2, ΔR1R3, ΔR2R3, ΔR1R2R3) were cloned into an ampicillin-resistant pET31b vector without the N-terminal signal peptide (aa 1-24) and with a C-terminal 6xHis-tag while the FapC wildtype (wt) was cloned into a kanamycin-resistant pET28a vector. The full amino acid sequences are included in Fig. S1. All proteins were expressed in *E. coli* and purified as described^*24, 66*^. Eluted protein fractions were immediately frozen in liquid nitrogen and stored at −80°C until further use.

### Purification of S6

For reasons of availability, we used double mutant of S6 rather than wt S6. Two Cys residues were inserted in the N- and C-terminal parts of the protein which show high mobility in the crystal structure and therefore are expected to have little, if any, effect on overall protein stability. Accordingly, a pET28(+) expression vector was produced by Genscript (Piscataway, NJ) to express the 101-amino acid (aa) S6 from *Thermus thermophilus* containing the mutations Met1Cys and Phe97Cys (S6^M1C, F97C^). The protein was expressed using a protocol based on the original procedure^*60*^. The plasmid was transformed into *E. coli* BL21(DE3) using single pulse electroporation at 1800V in an Electroporator 2510 (Eppendorf, Hamburg Germany). The transformed cells were incubated for 1 h at 37°C in Super Optimal with Catabolite repression medium, spread on an LB agar plate containing 50 μg/mL kanamycin and grown O/N at 37°C. A single colony was resuspended in ∼ 2 mL LB medium, spread on freshly prepared LB agar plates and grown O/N at 37°C after which the cells were resuspended in a few mL LB medium and used to inoculate 4 L LB medium. The cells were grown at 37°C at 180 rpm, induced with 1 mM IPTG at OD_600_ ∼ 0.8, grown for additional 4 h before harvested by centrifugation (4000 rpm, 4°C, 20 min). The cell pellet was dissolved in a few mL 50 mM Tris pH 7.4 containing 1 mM TCEP, 50 mg/L RNase, 50 mg/L DNase and a tablet of Roche cOmplete Mini Protease Inhibitor Cocktail, and stored at −80°C. After thawing, the cells were lysed by sonication, using a Q500 sonicator (Qsonica, Connecticut USA) set to ten 20 s pulses with 10 s pause at 20% intensity, using a 1.6 mm Microtip probe. The lysate was centrifuged (30,000 g, 4°C, 30 min) and the pellet discarded. Nucleic acid contaminants were removed by addition of 0.5% w/v polyethyleneimine and stirring for 20 min at 4°C after which the solution was centrifuged again. The protein content was precipitated from the supernatant with 70% (NH_4_)_2_SO_4_ and stirring for 1 h at 4°C. The solution was centrifuged (34,000 g, 45 min) and the supernatant discarded. The pellet was resuspended in MQ water with 1 mM TCEP and dialyzed against 50 mM Tris-HCl (pH 7.5) and 1 mM TCEP for 24 h. After filtration through a 0.22 μm filter, the solution was loaded onto a CM-Sepharose column coupled to an Äkta Pure system (GE Healthcare Life Sciences, Brøndby, Denmark). The sample was loaded on a column equilibrated with buffer A (50 mM Tris-HCL, 1 mM TCEP, pH 7.5) and run with 1 mL/min. The protein was eluted with a gradient of buffer B (buffer A + 1M NaCl) in 1 mL fractions. Based on SDS-PAGE, S6 was identified in the second major peak on the chromatogram (data not shown) and the S6-containing fractions were pooled and dialyzed against 50 mM Tris-HCL (pH 7.5) and 1 mM TCEP for 24 h. Protein concentration was determined using an ε_280_ = 12,700 M^-1^cm^-1^ and a molecular weight of 11.97 kDa on a NanoDrop 1000 (Thermo Scientific). S6 was aliquoted into 1 mL fractions of 1 mg/mL, frozen and lyophilized for later use.

### Circular dichroism spectroscopy

CD measurements were performed on a Chirascan-plus CD spectropolarimeter (Applied Photophysics, Leatherhead, UK). Lyophilized protein was weighed out on an analytical scale and dissolved directly in FA (or FM) of different concentrations. Both protein concentration and solution pH were measured afterwards. In control experiments, protein solutions were adjusted to similar pH values using HCl or NaOH. FA and FM absorb strongly below 240 nm, precluding the use of far-UV CD measurements. Protein denaturation was instead monitored in the near-UV (aromatic) CD region which reports on tertiary structure. All three proteins contain aromatic amino acids to different extents (HEWL: 6 Trp and 3 Tyr, S6: 1 Trp and 4 Tyr, Ubi: 1 Tyr). Denaturation was followed with 1 mg/mL protein at 290, 282 and 285 nm for HEWL, S6 and Ubi, respectively. The samples were transferred into a 3 mm quartz cuvette with a cap to avoid evaporation. Thermal scans were carried out between 5°C and 95°C with a scan rate of 1°C/min, step resolution of 0.1°C, 1 s time per point, and bandwidth of 2 nm. Data were fitted in KaleidaGraph using the following equation:

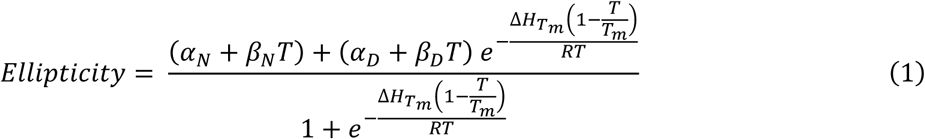

Here α_N_ and α_D_ are the ellipticity values of the native (N) and denatured (D) state, respectively, at 25°C and β_N_ and β_D_ describe the linear temperature dependence of the ellipticities of these two states. ΔH_*T*m_ is the enthalpy of denaturation at the midpoint of denaturation, *T*_m_. (NB: *T*_m_ is in units of Kelvin; we use the term *t*_m_ to refer to the value in Centigrade) This equation does not include specific heat capacity ΔC_p_ since this value cannot be determined with sufficient accuracy from a single thermal scan. However, by determining ΔC_p_ separately (see Results), the free energy, ΔG_T_ at 25°C can be determined using the following equation:

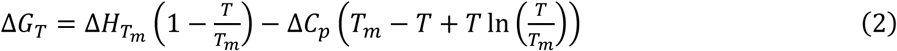

Using these ΔG values we calculate the fraction of natively folded protein (*f*_N_) as follows:

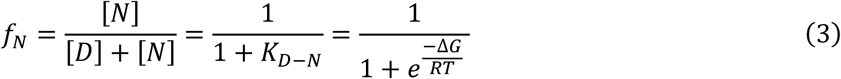

### Fluorescence spectroscopy

2 mg/mL protein stocks of lysozyme, S6 and ubiquitin were prepared from lyophilized material solubilized in MQ. The protein stocks were diluted into 1 mL samples with final concentrations of 1 mg/mL (lysozyme), 0.5 mg/mL (S6) or 0.25 mg/mL (ubiquitin) in FA concentrations ranging from 0-45% (v/v). Lysozyme was also treated with 10-90% (v/v) FM. The samples were incubated at RT for 30 mins before measurements to ensure that equilibrium was achieved. Fluorescence measurements were performed on a Cary Eclipse Fluorescence Spectrophotometer (Agilent) using a 10 mm quartz cuvette under isothermal conditions (25°C). Trp fluorescence was measured by excitation at 295 nm (5 nm slit) and the emission spectrum was recorded between 305 and 450 nm (2.5 nm slit), while combined Trp and Tyr fluorescence was recorded by excitation at 280 nm and emission at 295-450 nm. All spectra were recorded with 1 nm resolution and scanning speed of 0.5 nm/sec. Shift in Trp or Trp+Tyr emission intensity as function of FA or FM concentration was fitted to the following equation:

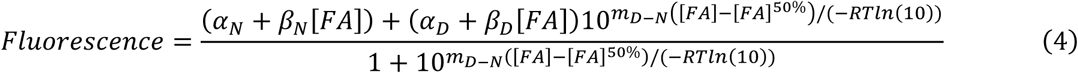

Here, analogously to eq. (2), α_N_ and α_D_ are fluorescence signal amplitude of the native and the denatured states, respectively, whereas β_N_ and β_D_ are the respective baseline slopes. [den]^50%^ is the denaturant concentration at which [N] and [D] are equal. The factor –*RT* ln(10) = −1.36 kcal/mol ensures that the *m*_D-N_ value reports on the linear dependence of Δ*G*_D-N_ (rather than ln *K*_D-N_) on [FA]. This equation was also used to monitor unfolding of 1 mg/ml S6 in 0-40% FA at RT using the near-UV CD signal at 282 nm.

### Fibril stability towards formic acid (FA)

Purified wt FapC and FapC mutants were desalted directly from elution buffer into pure MQ as described above. Concentration was measured by UV absorbance at 280 nm and the samples were fibrillated as above. Fibrils were spun down (13,500 rpm, 15 min) and the concentration of protein remaining in the supernatant was measured. This allows for calculation of the amount of pelleted protein fibrils. The fibrils were then resuspended in MQ to a concentration of 1 mg/mL (assuming all fibrils were dissolved into monomers) and aliquoted into eppendorf tubes. The fibrils were pelleted again and the supernatant discarded before resuspending them at 1 mg/mL in different concentrations of FA followed by 10 min incubation at room temperature. The samples were again centrifuged and 20 µL of each supernatant was lyophilized for ∼ 1 h. The lyophilized material was resuspended in 20 µL of MQ, mixed with reducing loading buffer (G Biosciences) and run on SDS-PAGE at 150 V for ∼ 70 min together with a prestained PageRuler protein ladder (Thermo Scientific). Gels were stained for 45 min in Coomassie Brilliant Blue (CBB) solution (1.2 mM CBB, 5% ethanol, 7% acetic acid) and destained O/N in destaining solution (5% ethanol, 5% acetic acid). Gels were quantified using Gel Analyzer 2010a^*78*^ and normalized to the band where fibrils were dissolved in 100% FA. We derive a predicted relationship between the fraction of dissolved protein and [FA] as follows. We assume a linear relationship between the free energy of unfolding in FA (Δ*G*_D-N_^FA^) and [FA]:

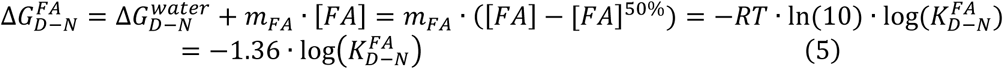

where [FA]^50%^ is the FA concentration where the fibrillar state (F) and the monomeric state (M) are equally stable, *i.e.* Δ*G*_D-N_^FA50%^ = 0 ↔ Δ*G*_D-N_^water^ = -*m*_FA_*[FA]^50%^ and operationally define *K*_D-N_^FA^ = [M]/[F] (analogous to the relationship between native and unfolded protein in conventional unfolding). This leads to the following expression for the percentage dissolved protein, *y*_*diss*_, as function of [FA]:

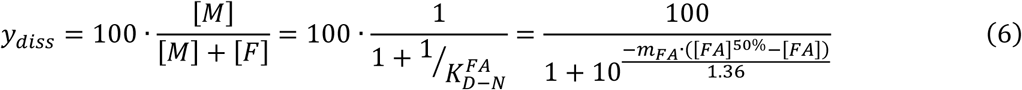

In this model, no protein dissolves in the absence of FA and all protein is dissolved at high [FA]. This is consistent with our observations.

### Fourier-transform infrared (FTIR) spectroscopy of the different fibrils

2.0 μL of fibrillated sample was dried onto an Attenuated Total Reflection (ATR) crystal with dry nitrogen on a Tensor 27 FTIR instrument (Bruker Optics, Billerica, MA). OPUS version 5.5 was used to process the data, including calculating atmospheric compensation, baseline subtraction, and second-derivative analysis. All spectra were made as accumulations of 68 scans with a resolution of 2 cm^-1^ in the range of 1000-3998 cm^-1^. Only the 1600-1700 cm^-1^ range, comprising information about secondary structure, is shown.

### Fibril stability towards SDS

Fibrils formed by wt FapC and FapC mutants were produced as follows: The FapC solution was desalted from elution buffer (8 M GdmCl, 50 mM Tris-HCl, 300 mM imidazole, pH 8) into 50 mM Tris-HCl, pH 7.4, in 0.5 mL fractions using a pre-equilibrated PD-10 desalting column (GE Healthcare) and concentration was measured on a NanoDrop using a theoretical extinction coefficient of 10095 M^-1^cm^-1^ calculated from the amino acid sequences (all FapC variants contain 1 Trp, 3 Tyr and 2 Cys) using ExPASy (https://web.expasy.org/protparam/). Protein solutions were then allowed to fibrillate O/N at 37°C with 900 rpm shaking in a table Eppendorf shaker (Thermo Scientific). After 24 h incubation, equal amounts of the fibrillated sample were aliquoted into 8 tubes and the fibrils were pelleted by centrifugation (13,500 rpm, 15 min) in a MicroStar12 table centrifuge (VWR) and the supernatants were removed and the concentration of un-fibrillated monomer was measured to determine the exact amount of fibrillated protein. Next, a non-reducing (58 mM SDS, 83 mM Tris-HCl, pH 6.8) and a reducing (58 mM SDS, 83 mM Tris-HCl, 1 mM TCEP, pH 6.8) SDS solution were prepared. The fibril samples were resuspended in 60 µL of either non-reducing or reducing SDS solution, yielding a final FapC concentration of 3.25 mg/mL. The samples were then incubated at 21°C, 50°C, 75°C or 95°C for 10 min. All samples were centrifuged as described previously and the supernatant of each sample was transferred to new tubes. Glycerol (10% (w/v)) and bromophenol blue (0.005% (w/v)) were added to the supernatants and the samples were analysed with SDS-PAGE. The intensity of the protein bands was quantified using ImageJ (https://imagej.net/). The percentage of solubilized FapC fibril was calculated for each sample by comparing their values to a dilution series of monomeric FapC protein of known concentrations.

## ACKNOWLEDGEMENTS

D.E.O. and L.F.B.C. are funded by Innovation Foundation Denmark (Grant 5188-00003B) through the Joint Programme on Neurodegenerative Diseases (aSynProtec). D.E.O. and J.S.N. are funded by the Independent Research Council Denmark | Technology and Production (Grant 9041-00123B) and the Lundbeck Foundation (Grant R276-2018-671). D.E.O. and T.S. are funded by the Independent Research Council Denmark | Natural Sciences (Grant 8021-00208B). We appreciate helpful discussions with Tom Record and Emily Zytkiewicz on *m*-values.

## AUTHOR CONTRIBUTIONS

L.C.: Formal analysis, Investigation, writing - original draft, writing-review and editing. J.S.N.: Formal analysis; investigation, writing-original draft. T.S.: Formal analysis, investigation. S.A.F.: Formal analysis, investigation. D.E.O.: Conceptualization, formal analysis, funding acquisition, methodology, project administration, supervision, writing - original draft, writing – review and editing.

## FOOTNOTES

## Abbreviations

aa: amino acid
ATR: Attenuated Total Reflection
CD: circular dichroism
cac: critical aggregation concentration
*E. coli*: *Escherichia coli*
FA: formic acid
fap: functional amyloid in *Pseudomonas*
FTIR: Fourier-transform infrared
GdmCl: guanidinium chloride
HEWL: hen egg white lysozyme
MQ: deionised water
MRE: mean residue ellipticity
O/N: overnight
RT: room temperature
SASA: solvent-accessible surface area
SDS: sodium dodecyl sulfate
wt: wildtype

